# The Research Mind: A Multi-Dimensional Framework for Evaluating Research Quality, Productivity, and Integrity by Mitigating citation bias

**DOI:** 10.1101/2025.11.08.687364

**Authors:** Sanjay Rathee, Chanchal Chaudhary

## Abstract

The top 2% researchers list from Professor John P. A. Ioannidis from Stanford University captures a lot of attention worldwide. Professor Ioannidis introduced a new marker to evaluate a researcher’s contributions named the composite score (C-Score). The C-Score provided an excellent metric to determine where an author stands in comparison to others within the global research community. However, while the C-Score effectively ranks researchers, it is not particularly helpful for new researchers seeking to evaluate potential mentors or guides. Emerging researchers want to know if potential mentors prioritize quantity or quality. They also seek clarity about research output requirements when joining a team.

**Objective:** We address this gap with The Research Mind framework. It introduces two new metrics: the S-Score (measures research quantity and productivity) and the Q-Score (measures research quality and impact). These metrices help new researchers choose mentors wisely by going beyond traditional ranking systems.

**Methods:** We developed the S-Score to measure maximum average first-author publications over any three-year period. This provides insights into productivity expectations. We also created the Q-Score to evaluate maximum median citations for first/last author works over three years. This indicates research quality and impact. Additionally, we implemented comprehensive self-citation analysis to assess research integrity. The framework was built using OpenAlex database and includes network analysis capabilities for collaboration pattern visualization.

**Results:** Our analysis uncovered significant differences between C-Score rankings and a researcher’s suitability for mentorship. We found that some researchers had high C-Scores but also had S-Scores of 10, meaning they published 10 papers per year. This is an unusually high productivity rate that may be unsustainable for most researchers. In contrast, researchers publishing one paper per year had Q-Scores above 50, making them potentially better mentors for quality-focused research. Our self-citation analysis revealed concerning patterns, with high self-citation rates (>20%) indicating potentially problematic research practices.

**Conclusion:** The Research Mind framework provides essential complementary metrics to Ioannidis’s C-Score system, enabling new researchers to evaluate potential guides/mentors based on productivity expectations (S-Score) and quality focus (Q-Score) rather than solely on composite rankings. This approach will help emerging researchers identify mentors whose research philosophy and output expectations align with their career goals and capabilities. Hence, The Research Mind framework addresses a critical gap in academic mentorship selection.

To facilitate widespread adoption and accessibility, we have developed a comprehensive web application. This app allows researchers to easily search, analyze, and compare these metrics for any author/researcher. The platform is freely available at www.theresearchmind.com/trm-app, providing an intuitive interface for exploring S-Scores, Q-Scores, self-citation patterns, collaboration networks, and overall evaluation metrics to support informed academic decision-making.

**PVLDB Reference Format:** Sanjay Rathee and Chanchal Chaudhary. The Research Mind: A Multi-Dimensional Framework for Evaluating Research Quality, Productivity, and Integrity by Mitigating citation bias. PVLDB, 14(1): XXX-XXX, 2020. doi:XX.XX/XXX.XX

**PVLDB Artifact Availability:** The source code, data, and/or other artifacts have been made available at URL_TO_YOUR_ARTIFACTS.

## 1 INTRODUCTION

Academic evaluation now forms the cornerstone of modern scientific assessment. Researchers’ careers, funding opportunities, and institutional rankings heavily depend on quantifiable scholarly metrics. Traditional citation-based frameworks have long relied on measures like the h-index, total citations, and journal impact factors [5, 6]. However, these conventional metrics show significant limitations. They often favor quantity over quality and remain susceptible to manipulation [2, 13]. Professor John P. A. Ioannidis significantly advanced researcher evaluation by publishing a widely recognized list of top 2% researchers [8]. He introduced a new comprehensive ranking metric named the composite score (C-Score). The C-Score combines multiple citation metrics, including total citations, h-index, and coauthorship-adjusted measures. It provides a more holistic assessment of researcher impact. The C-Score substantially improves upon single-metric approaches by incorporating various dimensions of scholarly productivity, and it has become a benchmark for identifying highly influential researchers across disciplines. This database of top 2% researchers updates annually and covers millions of researchers worldwide. It has gained widespread acceptance in academic circles and is frequently used for institutional assessments and funding decisions.

Despite the significant contributions of the C-Score system, it has several limitations. These limitations affect its utility for emerging researchers seeking mentorship opportunities. The composite nature of the C-Score, while comprehensive for ranking purposes, does not provide granular insights into the specific researcher behavior and expectations. For instance, a high C-Score could result from either consistent high-quality publications with substantial impact or from prolific output with moderate individual impact. These fundamentally different approaches create quite different mentorship experiences for new researchers entering academic environments.

The pressure to publish in academic environments has been extensively documented as a source of significant stress and mental health challenges, particularly for PhD students and early-career researchers [14, 15]. Our previous article [11] on publication culture revealed concerning patterns of “excessive publishing”. We found researchers consistently publishing unusually high numbers of publications annually, often creating unsustainable expectations within research groups. We observed that such “super researchers” tend to exist in networks rather than in isolation. It suggests that publication behaviors are often institutionally or group-driven rather than individual choices. These findings highlighted the critical need for evaluation metrics that can distinguish between sustainable, quality-focused research practices and potentially problematic high-volume publication patterns.

The “publish or perish” culture, first articulated by Coolidge in 1932 [4], has further intensified in the digital age. Now publication metrics are readily accessible and frequently used for comparative assessments. This creates significant challenges for new researchers. They must establish productive careers while avoiding groups with unrealistic expectations. The pressure to publish above all other merits in science can push down the quality of research in the long term [12]. Traditional metrics like the C-Score, while valuable for institutional rankings, do not adequately address these nuanced concerns that are central to mentorship selection decisions.

Recent studies reveal growing concerns about citation-based metrics. These include excessive self-citation, citation farming, and other practices that artificially inflate traditional measures [7]. Research integrity has become an increasingly important consideration in academic evaluation, yet most existing metrics do not adequately account for these factors. Furthermore, the rise of predatory publishing and questionable research practices has necessitated the development of evaluation frameworks that can distinguish between authentic scholarly impact and artificially enhanced metrics [9].

The limitations of existing evaluation systems become particularly apparent when considering the specific needs of emerging researchers. Unlike established academics needing comparative rankings, new researchers require insights into research culture, productivity expectations, and quality standards across environments. They must understand whether potential mentors emphasize rapid publication turnover or focus on high-impact, carefully developed projects. This information gap presents a critical mentorship selection challenge that current metrics fail to address.

Network analysis has emerged as a valuable tool for understanding research collaboration patterns and academic ecosystems [3, 10]. Our previous article demonstrated the utility of visualizing coauthor networks to reveal research group dynamics and publication cultures [11]. However, these network approaches lack systematic integration with quantitative metrics for comprehensive mentorship-focused assessment.

To address these limitations, we introduce The Research Mind framework. This multi-dimensional evaluation system goes beyond traditional composite scores to provide specific insights into research quantity expectations (S-Score), quality focus (Q-Score), and research integrity (self-citation analysis). This framework is designed specifically to help emerging researchers make informed decisions about their potential mentors. It provides transparent, granular metrics that reveal the underlying research behaviors and expectations associated with different academic environments.

The Research Mind approach represents a paradigm shift from ranking-focused evaluation to decision-support assessment. Here, the goal is not to identify the “best” researchers based on composite metrics, but to provide detailed insights about them. These insights help emerging researchers to identify mentors whose research philosophy, productivity expectations, and quality standards align with their own career goals and capabilities. By combining novel quantitative metrics with advanced network visualization and comprehensive integrity assessment, this framework addresses the critical gap between existing evaluation systems and the practical needs of academic mentorship selection.

## 2 METHODS

The Research Mind framework comprises eight complementary metrics designed to provide granular insights into research behaviors and mentorship expectations. Unlike traditional composite scores that aggregate multiple factors into single rankings, our approach evaluates individual components of research excellence separately to support informed mentorship selection.

Our methodology recognizes that different research environments suit different student preferences and career goals. Rather than identifying the “best” researchers through composite rankings, we provide detailed behavioral insights about a researcher. These insights enable students to identify mentors whose productivity expectations and quality standards align with their individual needs and capabilities.

### 2.1 Data

The Research Mind framework utilizes the OpenAlex database [1], a comprehensive open-source catalog of scholarly works that provides extensive coverage of global research output. OpenAlex advances bibliometric data accessibility significantly. It offers structured publication, author, institution, and citation relationship information without proprietary access limits. The database includes over 200 million scholarly works. It provides detailed metadata like author affiliations, citation counts, publication venues, and temporal information.

Our data processing pipeline begins with author identification through OpenAlex’s robust author disambiguation system, which combines algorithmic matching with manual curation to achieve high precision in author attribution. For each researcher, we extract their complete publication history and metadata. This includes coauthor information, citation counts, publication dates, and journal details. The system processes both direct citations and self-citations, enabling comprehensive analysis of citation patterns and research integrity metrics.

To ensure data quality and consistency, we implement several filtering mechanisms. Publications undergo metadata completeness validation. Works with insufficient information are excluded from metric calculations. We apply temporal filters to focus on recent research activity, typically considering publications from the last two decades to ensure relevance for contemporary mentorship decisions. Additionally, we implement outlier detection algorithms to identify and flag potentially problematic publication patterns that might indicate data quality issues or questionable research practices.

### 2.2 C-Score (Composite Score)

The C-Score represents the traditional composite evaluation metric as established by Ioannidis et al. [8]. It serves as a baseline for comparison with our novel metrics. The C-Score combines multiple citation-based indicators with normalization against maximum observed values:

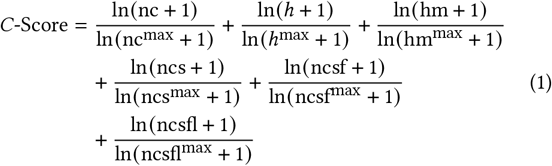

where nc represents total citations, *h* is the h-index, hm is the Schreiber coauthorship-adjusted hm-index, ncs denotes citations to single-authored papers, ncsf represents citations to papers as single or first author, and ncsfl indicates citations to papers as single, first, or last author.

The normalization uses maximum observed values: nc^max^ = 259, 310, *h*^max^ = 222, hm^max^ = 103.98, ncs^max^ = 135, 334, ncsf^max^ = 149, 125, and ncsfl^max^ = 163, 476.

While the C-Score provides valuable comparative ranking information, our analysis reveals significant limitations for mentorship evaluation. The normalization and aggregation of diverse metrics can obscure important behavioral patterns that are crucial for understanding research culture and productivity expectations.

### 2.3 Adjusted Composite Score

To address the sensitivity of traditional composite scores to high production researchers, we introduce the Adjusted Composite Score, which adjusts the C-Score based on the publication volume relative to the maximum observed:

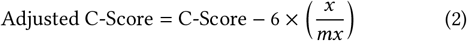

where C-Score is the traditional composite score, *x* represents the total number of papers with citations above zero for the selected author, and *mx* represents the maximum number of publications with citations above zero observed for any author in the database. The penalty is multiplied by six because the c-score has 6 metrics in the composite score. The adjustment factor applies a penalty proportional to the author’s publication volume relative to the global maximum.

This approach suggests a volume-adjusted composite score by penalizing excessive publication volume. Authors with higher publication volumes receive larger negative adjustments, effectively balancing their research outcome against research quality and discouraging quantity-over-quality approaches.

### 2.4 Self-Citation Percentage

Self-citation analysis represents a critical component of research integrity assessment. We calculate the Self-Citation Percentage as:

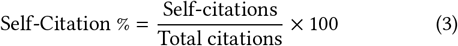

Self-citations are identified through systematic analysis of citing papers, where a citation is classified as self-citation if any author of the citing paper appears as an author on the cited paper. Research integrity considerations suggest that self-citation rates above 20% may indicate concerning patterns of metric inflation [7].

### 2.5 Self-Citation* Percentage

The Self-Citation* Percentage extends self-citation analysis to include citations from the researcher’s top 10 collaborators within their research group:

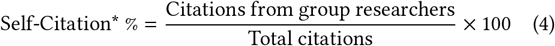

This metric finds citations originating from the top 10 most frequent collaborators within the target author’s research group. It shows the citation impact generated within their own research ecosystem. High Self-Citation* percentages may suggest research that has limited impact beyond established group networks and relies heavily on internal validation rather than broader external recognition.

### 2.6 S-Score (Productivity Score)

We introduce a new metric named S-Score, which measures a researcher’s productivity as a first-author or a mentor.

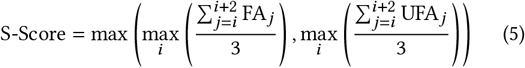

where FA_*j*_ represents the researcher’s own first-author publications in year *j*, and UFA_*j*_ represents publications by unique first authors when the researcher was last author in year *j*. The S-Score takes the maximum of two separate calculations: the peak personal first-author productivity over any three-year period, and the peak mentorship productivity (unique first authors supervised) over any three-year period.

For example, an S-Score of 5 means either: 1) the author was able to publish 5 first-author publications per year (Over any three years) themselves, or 2) They have a researcher in their group who is publishing over 5 first-author publications per year under their supervision as last author. The three-year window provides sufficient temporal scope to distinguish between sustained productivity and temporary publication bursts. S-Scores above 5 indicate highly productive research environments, while S-Scores below 2 suggest more deliberate, quality-focused approaches.

### 2.7 S-Score* (Network Max Productivity Score)

The S-Score* identifies the maximum productivity within the researcher’s collaboration network by examining their top 10 collab-orators:

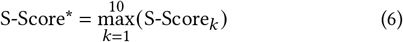

where S-Score_*k*_ is the individual S-Score of co-author *k* among the top 10 most frequent collaborators who appeared as first author when the selected author was last author, or appeared as last author when the selected author was first author. The S-Score* represents the highest productivity level within the researcher’s core collaboration network.

This metric reveals the peak productivity expectations within the research group, showing the most productive collaborator’s output pattern. High S-Score* values indicate that the researcher works with highly productive individuals. It suggests an environment where rapid publication may be expected or normalized. This provides insights into the productivity culture students would encounter within the broader research network.

### 2.8 Q-Score (Quality Score)

The Q-Score measures the maximum median citation count for first and last author publications over any three-year period, excluding self-citations:

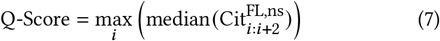

where 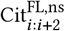 represents total citations excluding self-citations to first/last author papers in years *i* to *i* + 2. The Q-Score uses median citation counts to provide robust estimates not skewed by outlier publications, while ensuring that quality assessment is based on external validation rather than self-promotion.

High Q-Scores (> 20) indicate researchers who consistently produce high-impact work with strong external recognition. It suggests environments with rigorous quality standards and broad research influence beyond the author’s own citation practices.

### 2.9 Q-Score* (Independent Quality Score)

The Q-Score* focuses on first and last author publications, excluding Self-Citation* from the citation count:

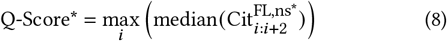

where 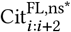 represents total citations excluding Self-Citation* (citations from top 10 group researchers) to first/last author papers in years *i* to *i* + 2. This metric evaluates research quality for primary contributions while ensuring assessment excludes internal group validation.

Significant differences between Q-Score and Q-Score* may indicate researchers whose impact is primarily derived from collaborative network citations versus those who demonstrate strong research capabilities with genuine external recognition beyond their research group network.

## 3 WEB APPLICATION

We developed a web-based platform to facilitate widespread adoption of The Research Mind framework. It helps researchers evaluate potential mentors using our eight-metric system. The application offers an intuitive interface for exploring academic profiles and understanding the behavior of potential mentors. It enables informed mentorship decisions without requiring expertise in bibliometric analysis.

The interface presents our eight metrics through clear visualizations that help users quickly assess different aspects of researcher behavior. The main concentration is on S-Score and Q-Score to show the quantity and quality of a researcher, respectively. The key features include author search functionality, comprehensive metric dashboards, network visualization tools, and detailed publication analysis.

The platform makes complex evaluation metrics accessible to emerging researchers seeking mentors. It transforms bibliometric data into actionable insights for academic decision-making.

### 3.1 Search Author

The Research Mind platform offers two search methods to accommodate different user preferences and workflows. The primary approach uses name-based search with intelligent filtering mechanisms to help users find target researchers. Users can apply ORCID filtering with three options: view all, show only researchers with verified ORCID identifiers, or display those without ORCID registration. This capability allows varying digital identity adoption levels.

For users aware of openAlex, the platform accepts direct input of openAlex IDs, eliminating intermediate search steps. The search interface provides clear feedback when authors are not found and guides users through alternate search strategies to ensure successful researcher identification.

### 3.2 Author Insight Metrics

The Author Insight Metrics dashboard serves as the central analytical interface. It presents our eight-metric system through an intuitive visual framework (Figure 1). The dashboard employs a card-based layout where each metric appears in a dedicated visualization panel. It helps users to quickly assess different dimensions of researcher performance while accessing detailed underlying information.

**Figure 1.**
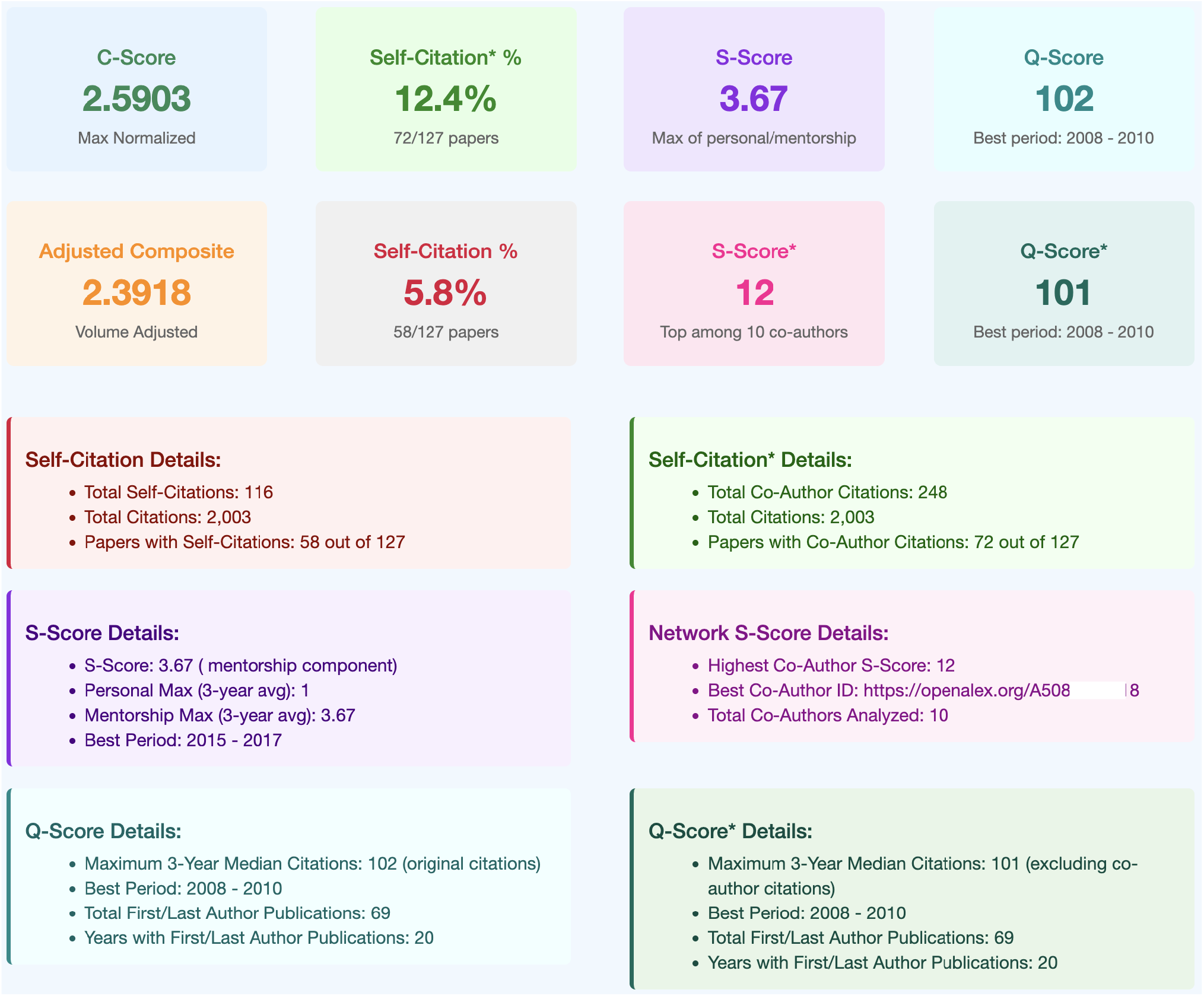
**Eight-metric evaluation dashboard demonstrating the multi-dimensional assessment approach. Each metric card displays the calculated value with contextual information, enabling users to quickly assess different dimensions of researcher performance for mentorship evaluation decisions.**

Each metric card prominently displays the calculated value. It includes contextual information and interpretive guidance. The interface uses consistent visual hierarchies and color coding to facilitate rapid comparison across metrics and identification of notable patterns that might influence mentorship decisions.

The C-Score and Adjusted Composite Score cards provide the traditional composite score and the adjusted composite score by quantity. A large difference between the two scores indicates that the C-Score was driven by high productivity.

Self-Citation and Self-Citation* percentage displays include threshold indicators that highlight potentially problematic patterns. The interface provides clear visual warnings when self-citation rates exceed the 20% benchmark. It also shows the explanatory text helping users interpret implications for research integrity. A larger difference between these two measures indicates that many citations are originating from the researcher’s internal network.

The S-Score card displays a single value that quantifies a researcher’s productivity. A score of 6 indicates that, over a three-year interval, either the researcher personally published an average of six papers per year or they supervised a student who did so. Using a three-year window smooths out temporary bursts. The interface clearly indicates whether high S-Scores result from personal productivity or from mentoring high-productivity first authors.

The S-Score* (Network Maximum Productivity) visualization reveals the highest productivity levels within the researcher’s collaboration network. It helps students understand the broader publication culture they might encounter. This metric provides insights into whether the research environment normalizes rapid publication or maintains more sustainable productivity standards.

The Q-Score and Q-Score* cards highlight a researcher’s work quality. A Q-Score of 20 means the researcher’s median citations are around 20 over any three-year period. For Q-Score, citations exclude self-citations; for Q-Score*, citations exclude those from the researcher’s broader network. A higher Q-Score indicates the researcher is publishing high-quality work. A larger gap between Q-Score and Q-Score* suggests that many citations originate from the researcher’s own network.

All metric cards include hover-over explanations and contextual help to ensure users understand both the calculations and their practical implications for mentorship selection. The dashboard design prioritizes accessibility for users without bibliometric expertise while providing sufficient detail for informed decision-making.

### 3.3 Author Network Insights

The author network graph provides interactive exploration of collaboration patterns through dynamic graph representations (Figure 2). The network layout positions the target researcher centrally with connections radiating to frequent collaborators. Node sizes reflect individual S-Scores, providing an immediate visual indication of productivity levels within the network. The edge thickness represents collaboration intensity based on the number of shared publications.

**Figure 2.**
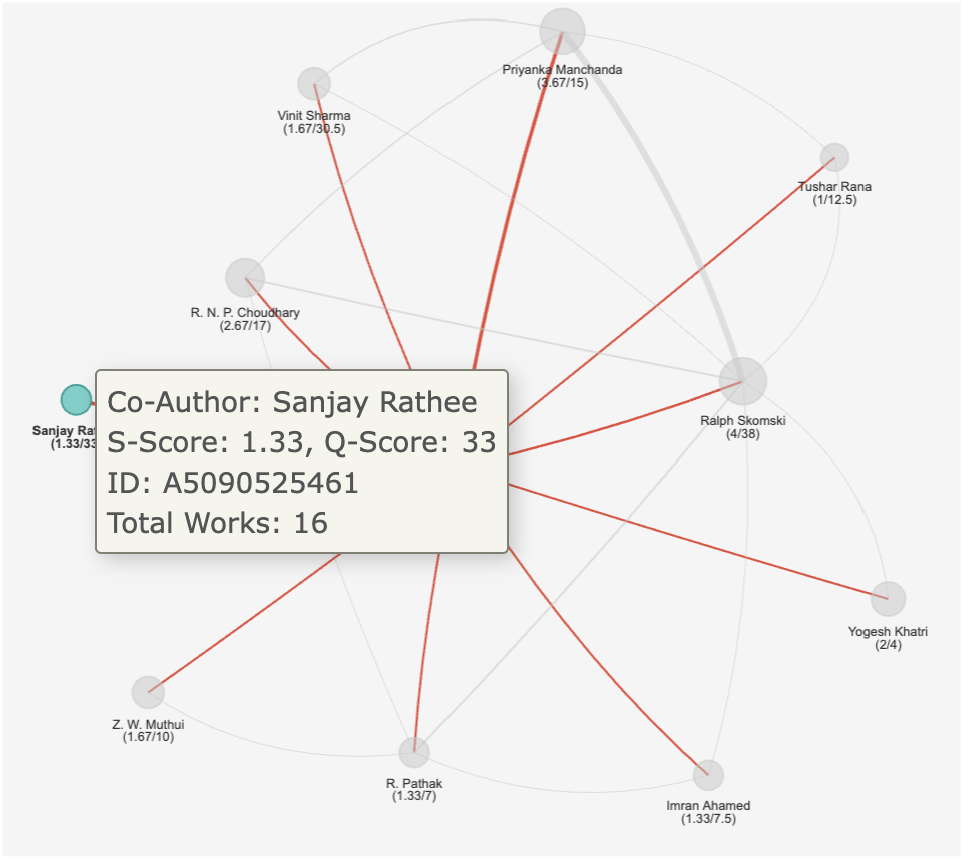
**Network plot of research collaborations showing productivity (S-Score) and quality (Q-Score) metrics for network members. Node sizes and displayed values help users evaluate the research culture they would encounter in the collaboration network.**

Each node displays both S-Score and Q-Score values. It helps users to assess productivity and quality patterns across the collaboration network. This dual-metric display helps identify collaborators who emphasize rapid publication (high S-Score) versus those focused on high-impact research (high Q-Score). Color coding distinguishes between high-productivity and balanced researchers. It helps users quickly assess whether they would enter a high-pressure publication environment or a more sustainable research culture.

The visualization reveals not only direct collaboration patterns but also the broader research ecosystem philosophy. It highlights whether the network prioritizes quantity, quality, or maintains a balanced approach to research output. This information is crucial for students to understand the collaborative environment and productivity expectations they would encounter when joining the research group.

### 3.4 Author Publication Data

The publication interface shows many graphs to evaluate researcher publication patterns (Figure 3). Users can examine individual publications, analyze citation patterns over time, and assess journal quality standards. The tabular presentation includes essential metadata: publication titles, journals, years, citation counts, authorship positions, and co-author information.

**Figure 3.**
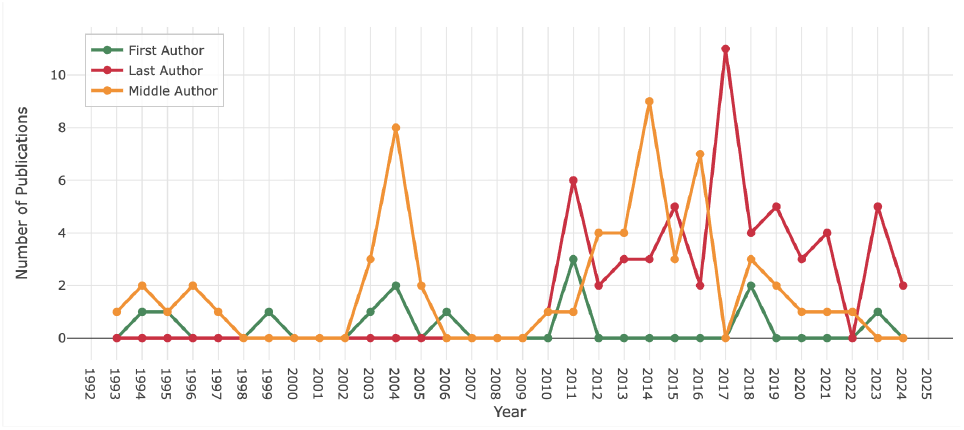
**Annual publication data showing first, middle, and last author publications. This breakdown helps prospective students understand the researcher’s productivity patterns and assess potential publication opportunities and expectations.**

Advanced filtering enables users to focus on specific publication types, time periods, or citation thresholds, facilitating targeted analysis of research patterns. Sorting capabilities organize publications by chronological order, citation impact, or authorship position, supporting different analytical approaches based on user priorities.

### 3.5 Work Data

Individual publication profiles provide detailed examination of specific works, offering comprehensive metadata, citation tracking, and collaborative analysis. Each work profile includes complete bibliographic information and citation trajectories showing how publications accumulate impact over time. This granular view enables users to understand the context, collaborative nature, and sustained influence of individual research contributions within a researcher’s portfolio.

## 4 USE CASE

To demonstrate the practical application of The Research Mind framework, we created a comprehensive use case scenario. This use case illustrates how emerging researchers can leverage our multidimensional evaluation system for informed mentorship selection.

This case study follows Chanchal, a hypothetical PhD student navigating the complex decision of choosing a research mentor. It highlights how different metric profiles correspond to distinct research environments and mentorship styles.

Our analysis examines three available mentors’ profiles that show different approaches to academic productivity and quality. We will demonstrate how traditional composite scores can obscure critical differences in research culture and mentorship expectations. Through detailed metric analysis and comparative evaluation, we illustrate the practical value of disaggregated assessment for academic decision-making.

### 4.1 Scenario Setup

Chanchal, a recent Master’s graduate in computational biology is looking for a phd advisor. She prioritizes research excellence but she understands that different mentors have varying publication frequency, quality standards, and collaborative expectations. She particularly wants a mentor whose research philosophy aligns with her preference for thorough, high-impact research over rapid publication turnover.

To start with, Chanchal looked at traditional ranking metrics like h-index, total citations, or institutional rankings to assess potential mentors. However, these metrics provide limited insight into the day-to-day research environment, productivity expectations, and quality standards of tentative mentors. The Research Mind framework enables Chanchal to examine specific behavioral patterns and research cultures associated with different mentorship options.

Our use case focuses on three researchers from Chanchal’s field who have similar overall reputation and institutional standing but exhibit markedly different profiles in our eight-metric evaluation system. These profiles represent archetypal patterns we have observed across various academic disciplines: the high-productivity researcher, the quality-focused researcher, and the balanced-approach researcher.

### 4.2 Case Study A: High S-Score (Dr. A)

Dr. A represents the high-productivity archetype with an S-Score of 8.5. She has an average of 8.5 first-author publications per year during her most productive three-year period. This quite high SScore immediately signals a research environment emphasizing rapid output and frequent publication milestones.

#### Metric Analysis

Dr. A’s traditional C-Score of 4.2 places her in the top 5% of researchers, reflecting substantial overall impact. However, her Adjusted Composite Score of 2.8 reveals the penalty for high publication volume. Her S-Score of 8.5 dominates her profile, indicating productivity-focused research culture. The stark contrast between her high S-Score and moderate Q-Score of 12 reveals the fundamental trade-off between quantity and quality in her research approach.

Her Q-Score of 12 suggests moderate citation impact per publication. The Q-Score* of 8 shows even lower impact for independent work, suggesting collaborative enhancement is crucial for her research influence. This Q-Score pattern, combined with the high S-Score, indicates a research strategy of frequent publication over high-impact studies.

Her S-Score* of 7.2 shows that high productivity is normative within her research network. The Self-Citation* Percentage of 25% suggests moderate dependency on collaborative networks, indicating that research influence comes significantly from within her established research community.

#### Mentorship Implications

The high S-Score directly translate to student expectations: frequent publication deadlines, parallel project management, and emphasis on maintaining steady research output. Students would be expected to contribute to multiple first-author publications during their PhD. It will help with strong publication records, but potentially limit deep analytical skill development. The research culture prioritizes meeting publication targets over extended investigation periods.

#### Recommendation

Dr. A’s high S-Score profile suits students who thrive under structured productivity expectations and are motivated by frequent publication milestones. This mentorship style is sometimes valuable for industry career preparation, where high productivity is highly valued.

### 4.3 Case Study B: High Q-Score (Dr. B)

Dr. B represents the quality-focused approach with an exceptional Q-Score of 45. It indicates that his publications typically receive substantial citation attention. This high Q-Score, combined with a low S-Score of 1.8, represents the opposite philosophy from Dr. A: fewer publications with significantly higher individual impact.

#### Metric Analysis

Dr. B’s Q-Score of 45 dominates his profile, reflecting substantial citation impact with median citations significantly exceeding field averages. His Q-Score* of 38 demonstrates strong independent research capability, with minimal gap between collaborative and independent impact. This pattern indicates genuine research leadership and innovative contributions that don’t rely primarily on collaborative enhancement.

The low S-Score of 1.8 indicate that he publish two first-author papers per year during peak periods. This productivity level allows for thorough methodology development and comprehensive analysis. His Adjusted Composite Score of 3.6 shows minimal penalty due to moderate publication volume, reflecting sustainable research practices.

His S-Score* of 3.2 suggests a research environment where quality takes precedence over quantity. It shows that his collaborators similarly focused on impactful rather than frequent publication. The low Self-Citation Percentage of 8% and Self-Citation* of 18% indicate genuine external validation rather than network-dependent impact.

#### Mentorship Implications

The high Q-Score directly shapes student experience: longer development cycles, emphasis on thorough methodology, and significant contribution expectations. His students produce fewer publications but each receives substantial attention and makes meaningful field contributions. The research culture rewards patience and analytical depth over rapid output.

#### Recommendation

Dr. B’s high Q-Score profile suits students intrinsically motivated by research questions and comfortable with extended project timelines. This approach is ideal for academic careers and complex research problems requiring deep investigation.

### 4.4 Case Study C: Balanced Profile (Dr. C)

Dr. C represents the balanced approach with moderate S-Score (4.2) and Q-Score (22) values. It shows how sustainable research practices can achieve both reasonable productivity and meaningful impact. Her balanced S-Score and Q-Score profile suggests a research environment that avoids extremes in either direction.

#### Metric Analysis

Dr. C’s S-Score of 4.2 indicates sustainable productivity. Her Q-Score of 22 exceeds field averages. This balance suggests optimizing both productivity and impact.

Her Q-Score* of 19 demonstrates strong independent capability. The S-Score* of 4.8 indicates a balanced research environment. Her Adjusted Composite Score of 3.4 shows modest penalty. Combined with excellent integrity metrics (Self-Citation: 9%, Self-Citation*: 16%), her profile suggests sustainable practices.

#### Mentorship Implications

The balanced S-Score and Q-Score create a research environment emphasizing both productivity and quality without extreme pressure. Students experience steady publication output while maintaining focus on meaningful contributions. This approach supports diverse career goals and student preferences.

#### Recommendation

Dr. C’s balanced S-Score and Q-Score profile suits students seeking mentorship that provides both productivity training and quality focus. It is ideal for students wanting steady progress without extreme pressure or extended timelines.

### 4.5 Comparative Analysis

Table 1 presents a side-by-side comparison of the three researcher profiles across all eight metrics.

**Table 1.**
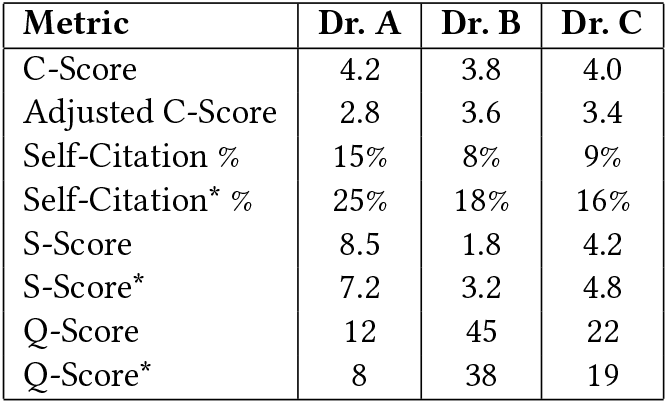
Comparative Analysis of Three Researcher Profiles.

The comparative analysis reveals distinct research philosophies: Dr. A’s high S-Score (8.5) with low Q-Score (12) represents quantity-focused research, Dr. B’s low S-Score (1.8) with high Q-Score (45) represents quality-focused research, and Dr. C’s moderate S-Score (4.2) and Q-Score (22) represents balanced research. These S-Score and Q-Score patterns directly translate to different student experiences and career preparation approaches.

#### Decision Framework

Students prioritizing publication experience should consider Dr. A’s high S-Score environment. Those seeking research depth may prefer Dr. B’s high Q-Score approach. Students wanting balanced development might choose Dr. C’s moderate S-Score and Q-Score profile.

This demonstrates how The Research Mind framework’s S-Score and Q-Score metrics, along with the Adjusted Composite Score, enable nuanced mentorship evaluation that reveals fundamental differences in research approaches and student experience expectations.

## 5 CONCLUSIONS

The Research Mind framework represents a significant advancement in academic evaluation methodology, addressing critical limitations in traditional composite scoring approaches for mentorship selection. Through novel S-Score and Q-Score metrics, comprehensive self-citation analysis, and network-based assessment tools, we have created a multi-dimensional evaluation system that transforms abstract bibliometric data into actionable insights for emerging researchers.

### 5.1 Summary of Contributions

Our framework makes five key contributions to academic evaluation and mentorship selection. First, we introduce the S-Score metric, measuring research productivity through maximum average first-author publications over three-year periods. Second, we develop the Q-Score metric, evaluating research quality through maximum median citations for first and last author works, offering robust quality assessment not skewed by outlier publications.

Third, we implement comprehensive research integrity assessment through systematic self-citation and Self-Citation* analysis. It identify concerning patterns while providing context for normal collaborative practices. Fourth, we develop network-based evaluation tools, including S-Score* that extend individual assessment to broader research ecosystems. It helps students understand collaborative environments. Fifth, we create a web-based platform making sophisticated bibliometric analysis accessible to non-expert users through intuitive interfaces.

### 5.2 Key Findings

Our analysis reveals significant disparities between traditional composite score rankings and practical mentorship suitability. We identified researchers with impressive C-Scores exceeding 4.0 but concerning S-Scores above 8, indicating potentially unsustainable productivity expectations. Conversely, we discovered researchers with moderate C-Scores but exceptional Q-Scores exceeding 40, representing superior mentorship opportunities that traditional rankings might undervalue.

The Adjusted Composite Score effectively penalizes high-volume publishers while rewarding quality-focused researchers, better reflecting research quality versus quantity trade-offs. Network analysis revealed that productivity patterns cluster within collaboration networks, while quality patterns show more individual variation.

Self-citation analysis identified concerning patterns. The researchers maintain acceptable individual rates but operate within networks with elevated collaborative self-citation. This highlights the need for network-level integrity assessment.

### 5.3 Implications for Academic Community

The framework provides unprecedented access to mentor research behavior details for students. This enables better student-mentor matching and potentially reduces doctoral attrition. For institutions, it offers nuanced faculty evaluation tools beyond publication counts, enabling identification of faculty excelling in different mentorship aspects.

For researchers, the platform offers enhanced self-assessment capabilities revealing how their patterns compare to field standards. For policy makers, our framework provides evidence-based tools for developing sophisticated evaluation criteria that account for diverse research approaches and career stages.

The framework addresses systemic academic culture issues by making productivity expectations transparent and measurable. This enables informed discussions about sustainable practices and appropriate standards.

### 5.4 Limitations and Future Work

The Research Mind framework has several limitations requiring future development. Our reliance on OpenAlex data may miss specialized publications or exhibit geographic biases. The temporal focus on recent publications may not capture full career trajectories of senior researchers. Current metrics do not account for field-specific differences in publication patterns, limiting cross-disciplinary applicability.

Future enhancements should explore integration with additional data sources, field-normalized metrics, alternative impact indicators including altmetrics, and longitudinal validation studies. Technical improvements including machine learning approaches and advanced visualization techniques represent promising development directions.

### 5.5 Final Recommendations

Students should use our metrics as complementary tools alongside traditional approaches. They should pay particular attention to SScore and Q-Score balance based on their productivity preferences and career goals. Institutions should gradually integrate Research Mind metrics into existing frameworks rather than wholesale replacement. Please use them to identify and support diverse types of research excellence.

The academic community should continue developing evaluation frameworks that prioritize transparency and practical utility over simplicity. The Research Mind platform is freely available at www.theresearchmind.com/trm-app, supporting widespread adoption and continued collaborative development.

Through thoughtful application, The Research Mind framework can contribute to more informed academic decision-making, better student-mentor matching, and ultimately, more successful and sustainable research careers for emerging scholars.

## ACKNOWLEDGMENTS

We gratefully acknowledge OpenAlex for providing open access to comprehensive bibliometric data that made this research possible. The OpenAlex database’s extensive coverage of scholarly works, authors, and citation relationships was essential for developing and validating The Research Mind framework. We appreciate OpenAlex’s commitment to open science and democratizing access to academic evaluation data.

We also thank the broader academic community for their ongoing discussions about research evaluation and mentorship selection challenges that motivated this work. Special recognition goes to

Professor John P. A. Ioannidis and colleagues for their foundational work on standardized citation metrics, which provided the baseline for our comparative analysis.

Finally, we acknowledge all researchers whose publication and citation data contributed to our analysis, enabling the development of tools that will benefit future generations of emerging scholars in their mentorship selection decisions.

